# NeatSeq-Flow: A Lightweight High-Throughput Sequencing Workflow Platform for Non-Programmers and Programmers Alike

**DOI:** 10.1101/173005

**Authors:** Menachem Sklarz, Liron Levin, Michal Gordon, Vered Chalifa-Caspi

## Abstract

Biologists often find it necessary to execute bioinformatic workflows (WFs) as part of their research. However, operation of most WF-management platforms requires at least some programming expertise. Here we describe NeatSeq-Flow, a platform that enables users with no programming knowledge to design and execute complex high-throughput sequencing WFs on their own computer or computer cluster. Workflows are composed of modules. NeatSeq-Flow provides a large compendium of pre-built modules as well as a generic module. Advanced users can also generate custom-made, sophisticated modules using templates and only basic Python commands. Modules and WFs are easily shareable. To execute a WF, through either the graphical user interface or the command line, users need to only specify modules’ order and parameters (workflow design) and input file locations (sample information). WF execution is parallelized on both samples and analysis steps, and progress can be tracked in real time. Results are obtained in a neat directory structure, along with a self-sustaining WF backup for reproducibility. NeatSeq-Flow operates by shell-script generation, allowing full transparency of the WF process. NeatSe q-Flow supports CONDA for easy installation and portability of entire environments. All these features make NeatSeq-Flow an easy-to-use WF platform without compromising flexibility, reproducibility, transparency and efficiency.

**Availability:** http://neatseq-flow.readthedocs.io/en/latest/

## Introduction

Modern biological experiments involving High-Throughput Sequencing (HTS) produce large amounts of data, which scientists must analyze in order to reach the kernel of information of interest. Usually, analysis of the data is composed of several operations, each of which consists of calling a program with inputs, receiving the outputs and passing them on to the next step. Often, the analysis is parallelized on multiple processing units (CPUs) or cluster nodes, thus saving execution time. The bioinformatician will typically write short shell scripts that execute the different operations and send them sequentially to a computer cluster job scheduler for execution on distributed nodes.

Creating and executing these script-based workflows (WFs) is time consuming and error prone, especially when considering projects with hundreds or thousands of samples, with many steps and plenty of intermediate files, or when the same analysis has to be repeated with different combinations of programs and parameters.

To address these and other issues, many commendable efforts have been made to create platforms for automating execution of such WFs (for examples, Refs. 1-6), a review of which was published a couple of years ago (7).

Most of the available WF platforms fall into two main categories: systems using a graphical user interface (GUI, e.g. Galaxy (8)) and command-line based systems (e.g. Nextflow (1), Snakemake (2) and SUSHI (3)). While intended for scientists with no programming experience, GUI-based systems usually have limited flexibility and transparency, and they often do require programming expertise in order to assimilate new tools or to perform complex WFs. On the other hand, command-line based systems are much more flexible and enable tailor made WF designs. However, command-line based systems require programming (e.g. in Groovy, Python or Ruby) even at the design stage of the WFs and are intended for dedicated, expert bioinformaticians.

We have developed NeatSeq-Flow, a lightweight, easy to use, yet powerful WF platform, which offers the advantages of the two worlds presented above. NeatSeq-Flow can be executed either from the command-line or using a dedicated GUI. It is easy to use for non-programmers and programmers alike, and in the same time it provides great flexibility and power.

NeatSeq-Flow as well as it’s GUI are written in Python and are easily installed, optionally using the CONDA package, dependency and environment manager (https://conda.io). NeatSeq-Flow is modular, can use existing as well as newly devised modules, and can execute both publicly-available and in-house programs. Ready-to-use workflows are available for common Bioinformatics analyses such as RNA-Seq, ChIP-Seq, variant calling, shotgun and amplicon metagenomics (https://neatseq-flow.readthedocs.io/projects/neatseq-flow-modules/en/latest/#neatseq-flow-workflows).

## Main Advantages

NeatSeq-Flow WFs are conceptually based on three elements: sample information (files’ physical location and type), modules and WF design. This setup enables advantages such as the use of the same WF design on different sets of samples as well as using different WFs on the same set of samples, all of this without changing the individual elements. Moreover, this independency of elements makes them easily shared between users and eventually forms a repository of WFs and modules ready to be used on new sample sets or to be re-edited to form new types of WFs and modules.

The three elements of a WF contain all the information required for its reproduction and are therefore stored by NeatSeq-Flow as a self-sufficient backup for this purpose. Execution of the WF is fully under the user’s control, using easily understood shell scripts generated by NeatSeq-Flow. These scripts also contain directives enabling parallelization and ensuring sequential execution. In addition, when executing WFs on a computer cluster the user can also determine to which node a step will be sent according to the step requirements such as the amount of memory and number of CPUs or by specifying a specific node name.

HTS analyses typically produce numerous intermediate and final files. NeatSeq-Flow automatically determines and manages the location of these files, and handles their transfer between WF steps. By the end of a WF execution, all files are neatly organized in an intuitive directory structure.

Designing and running NeatSeq-Flow WFs using existing modules does not require any programming knowledge and with the use of the included “generic module” most Linux-based programs having command-line arguments are also covered, making NeatSeq-Flow accessible to a wide variety of users.

NeatSeq-Flow is designed to be used locally, either on a single computer or on a computer cluster. Optionally, the user may also install NeatSeq-Flow GUI and use it locally. Thus, all analyses can be done within firewalls, and no trafficking of big data to remote servers is required.

Finally, NeatSeq-Flow supports the use of CONDA for easy installation of NeatSeq-Flow with most of its dependent HTS analysis programs. The use of CONDA environments does not require “superuser” privileges (“sudo”) and helps save time and effort in setting up a WF to work on the user’s own computer system. Moreover, in complex WFs CONDA relieves the user from the need to deal with interdependency issues among the installed programs. Most importantly, CONDA facilitates sharing of WFs by enabling delivery of entire environments for HTS analyses.

### Description of NeatSeq-Flow

NeatSeq-Flow can create and execute WFs on any set of samples and operations. A schematic diagram of NeatSeq-Flow and a detailed example are provided in Figs. 1 and S1.

**Figure 1.**
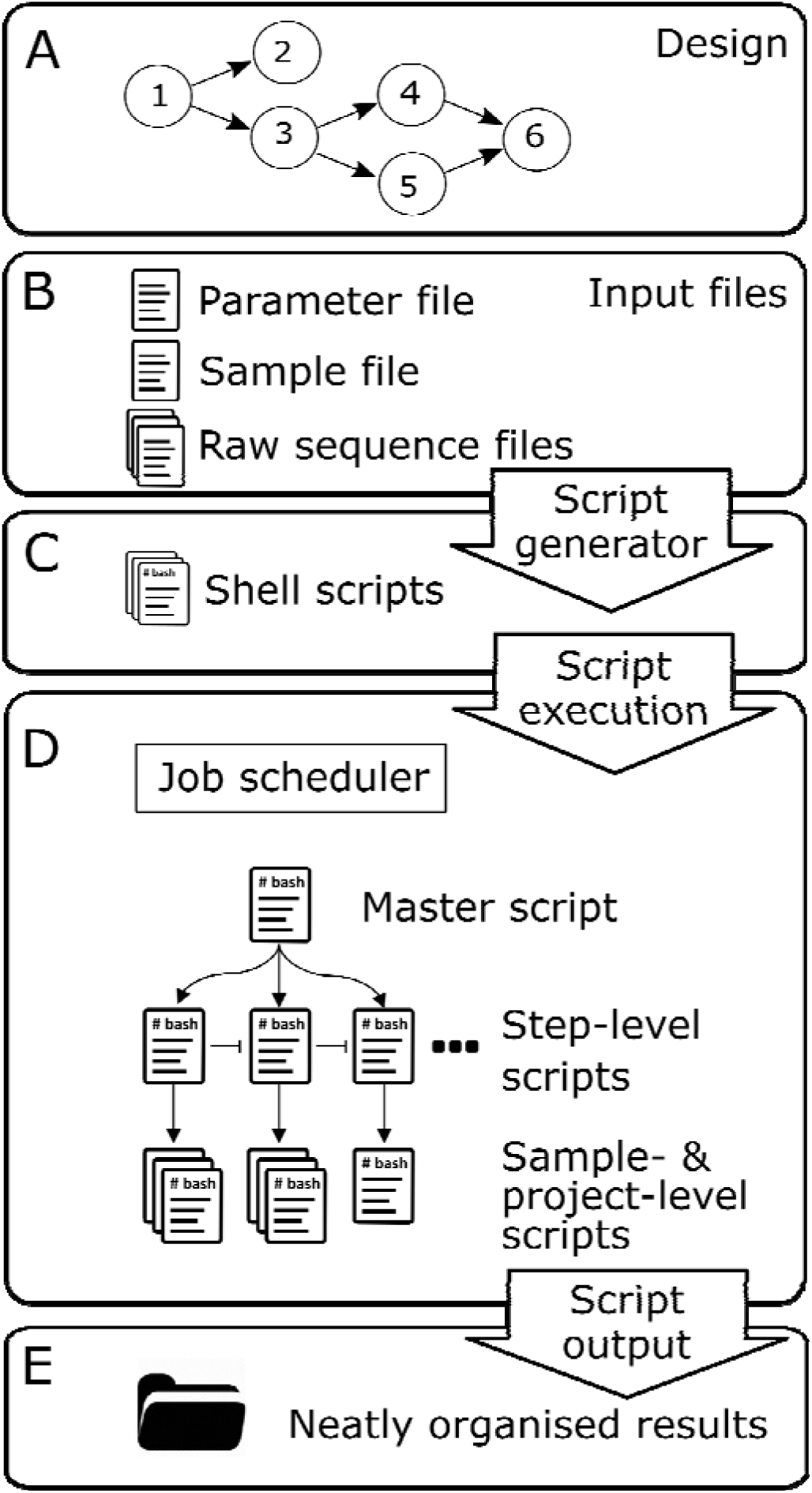
Outline of workflow execution with NeatSeq-Flow. **A.** A conceptual design of the WF takes on the form of a directed acyclic graph where nodes represent steps, arrows represent interdependencies between steps, and convergence (e.g. step 6) represents a step which is dependent on several previous steps. **B.** Based on the WF design, the user creates sample- and parameter-definition files, and provides the input files. **C.** NeatSeq-Flow Script generator is executed, creating a set of structured shell scripts. **D.** The shell scripts are typically executed on a computer cluster. Step dependencies are maintained through directives within the shell scripts. **E.** Script outputs, WF log and other accompanying files are neatly organized in a directory structure.

NeatSeq-Flow operations are implemented as modules, where each module is a wrapper for a program. A program could be anything executable from the Linux command-line, from a simple script to a complex software tool, either publicly available (e.g. Trinity (9), BWA (10), Bowtie (11), QIIME2 (12)) or an in-house program. A list of pre-built modules is available at NeatSeq-Flow module and workflow repository, http://neatseq-flow.readthedocs.io/projects/neatseq-flow-modules/en/latest/. Creation of new modules requires basic Python programming knowledge and is easily achieved using a provided template (Fig. S2). In addition, NeatSeq-Flow includes a generic module which can execute any Linux-based program having command-line arguments (in conventional formats). Usage of the generic module does not require any programming knowledge and may thus enable non-programmers to make full use of NeatSeq-Flow for running their own set of programs even if they are not available in NeatSeq-Flow module repository.

The order of the operations, i.e. the WF, is specified in a user-provided “parameter file” (see an example in Fig S1C). The parameter file may either be created manually using a text editor, or through NeatSeq-Flow GUI (See animated demonstration at https://github.com/bioinfo-core-BGU/NeatSeq-Flow-GUI). The WF is composed of steps, where each step calls a module. A certain module may be called by several distinct steps, e.g. each time with different parameters. For each step, the user defines which previous step(s) need to be completed before the current step executes, thus imposing “step dependencies”. The flow of steps may be perceived as a directed acyclic graph (Fig. 1A and see example in Fig. S1A), meaning that a step may be preceded either by a single step (e.g. Bowtie2 (13) precedes Samtools (14) in Fig S1A), or by a convergence of several steps (e.g. MultiQC MultiQC (15) in Fig S1A). Typically, steps are implemented at one of two levels: per sample (e.g. alignment to a reference) or per project (e.g. *de novo* assembly of the reads from all samples). Finally, for each step the user may define the module’s parameters (e.g. the use of the mem algorithm at the BWA mapper module in Fig. S1C), the program’s parameters (e.g. –B for mismatch penalty in BWA mem in Fig. S1C) and, optionally, step-specific cluster parameters (e.g. use nodes with certain memory/CPU requirements or run on specific node name(s)).

The set of raw input files (e.g. FASTQ and FASTA files) can be placed in a directory or directories of the user’s choice, and their location(s) and sample attributions should be defined in a “sample file” (see example in Fig. S1B). The “sample file” may be created manually using a text editor or through NeatSeq-Flow GUI. From this point onwards, the user is relieved from the need to know or manage the locations of intermediate or final files, or to transfer files between WF steps. WF output file locations are determined by NeatSeq-Flow such that they are neatly organized in an intuitive directory structure (Fig. S1E).

Once the user provides NeatSeq-Flow with sample and parameter files, NeatSeq-Flow creates a hierarchy of shell scripts (Fig 1C,D and Fig S1E): a “master script” that calls all step-level scripts; step level scripts that call all sample- or project-level scripts; and sample- and/or project-level scripts that call the relevant programs. The latter shell scripts contain the code for executing the programs, including input and output file locations, user-defined parameters and dependency directives. All scripts are stored in a neat directory structure (see example in Fig. S1E).

Execution of the WF takes place by running the WF’s master shell script (Fig. 1D). Step dependencies encoded in the shell scripts ensure the correct order of step execution. Parallelization is both sample-wise as well as step-wise for steps that are on independent branches of the WF (e.g. running several mapping programs as in Fig. S1A or running the same program with different parameter sets). The user may choose to execute only part of the WF; only a certain step; or even only a certain step on a certain sample, by executing the relevant shell script(s) from the script hierarchy. During WF execution, NeatSeq-Flow “Terminal Monitor” may be used to follow the WF progress in real time and to alert for execution errors (Fig S1D).

The WF output files are neatly organized in the “data” directory by module, step and sample (see example in Fig. S1E), making it easy to locate required information. Additionally, execution start and end times as well as maximum memory requirements are written to a log file. Debugging is facilitated by storing STDERR and STDOUT of the shell scripts in dedicated directories. All WF elements necessary for its execution, i.e. its parameter file, sample file and used modules, are copied into a dedicated backup directory. This enables reproducing the WF at any time in the future. Needless to say, the shell scripts themselves, together with the sample and parameter files, constitute the ultimate documentation for the WF performed. Sharing WFs is facilitated by a shared repository of modules and parameter files (http://neatseq-flow.readthedocs.io/projects/neatseq-flow-modules/en/latest/).

### Implementation

NeatSeq-Flow script generator, NeatSeq-Flow GUI and NeatSeq-Flow modules are written in Python. The parameter file uses the intuitive YAML format. Program paths (e.g. physical location of Bowtie2 executable) are specified by the user at the top of the parameter file and are easily edited, thus ensuring portability of NeatSeq-Flow and its modules across different computers (Fig. S1C). Input and output file paths of WF steps are determined “on the fly” by the script generator (see below), and are not hard coded in NeatSeq-Flow nor in the parameter file. This concept enables the modules and parameter files to be independent of actual file locations and therefore shareable.

Step dependencies are implemented as follows: *In the parameter file*, the user specifies for each step, which other step(s), called “base step(s)”, must precede it (Fig. S1C); *During execution of the script generator*, for each step, the script generator writes a directive into the relevant shell script to hold the execution of the current step’s program until the base step’s program terminates.

Information sharing between WF steps is implemented as follows: each module contains a definition of the required input file types and the expected output file types, e.g. for BLAST, the module defines FASTA and BLAST-database as inputs and BLAST result file as output. The output file locations are determined by the script generator for each step and stored in an internal data structure which is then passed on to the next step in the WF. In turn, each step can search the data structure for its required input file types (for more details see Fig S3). This design enables great flexibility for the user to thread together steps, with the only requirement being that for each step its required input file type(s) were created by at least one of its predecessor step(s).

The generic module does not contain a definition of input and output file types, therefore in steps that use a generic module, the user has to specify the input and output file types in the parameter file. An example of calling the generic module is provided in Fig. S4, and a full specification is available in NeatSeq-Flow documentation (http://neatseq-flow.readthedocs.io/projects/neatseq-flow-modules/en/latest/Module_docs/GenericModule.html)

To summarize, the script generator generates the shell scripts as follows: for each step it (1) writes a directive into the relevant shell script, to hold the execution of the current step’s program until the base step’s program terminates; (2) checks that the input file type(s) of the current step’s module are compatible with the output file types of its predecessors step(s); (3) constructs unique file paths for the outputs of the current step and stores them in the internal data structure as appropriate types; (4) retrieves from the data structure the file paths of its required input files, generated by previous steps; (5) constructs the shell command for calling this step’s program with the input and output file paths, and writes the command in the relevant shell script.

## Conclusions and Future Perspective

NeatSeq-Flow allows the user to execute diverse and extensive HTS analyses on computer clusters, while avoiding the tedious task of composing numerous error free shell scripts. Execution of the actual WF is controlled by information depicted in the shell scripts produced by NeatSeq-Flow, while the user has the freedom to choose which steps and which samples to execute. A WF in NeatSeq-Flow is defined by sample and parameter files and together with the modules used they ensure clear documentation and reproducibility. Furthermore, once the shell scripts are produced by NeatSeq-Flow, they plainly reveal all the operations applied to the data, with nothing “hidden behind the scenes”. NeatSeq-Flow is written in plain Python, such that adding new modules to the software is a straightforward process. A generic module is also provided, enabling calling programs directly, without pre-built modules. Accordingly, NeatSeq-Flow may easily be extended to include new protocols and software packages. It is our hope that the community of users will contribute additional modules as well as dedicated WF designs to the public. Programmers and non-programmers alike may benefit from easy WF design and execution of NeatSeq-Flow, through either the command-line or the GUI. NeatSeq-Flow is in constant use by our group for a multitude of analysis procedures, and has proven to be priceless in time saving and error reduction. NeatSeq-Flow is under continuous agile development and improvement. NeatSeq-Flow is general-purpose and may easily be adjusted to work on different types of analyses other than HTS.

## Supporting information

## Acknowledgements

This research used the High Performance Computing Facility at Ben-Gurion University. Conflict of Interest: none declared.

---

**Figure S1. Example of NeatSeq-Flow workflow: A.** Graphical presentation **B.** Sample file **C**. Parameter file **D**. Terminal Monitor **E**. Output directory structure

**Figure S2. Template for a new Module: A.** Sample-level module template **B.** Project-level module template

**Figure S3. Implementation of managing file transfer between steps**

**Figure S4. Example of usage and implementation of the generic module**

