## Supplementary material for "NeatSeq-Flow: A Lightweight High-Throughput Sequencing Workflow Platform for Non-Programmers and Programmers Alike"

**Availability**: <http://neatseq-flow.readthedocs.io/en/latest/>

**Contact**:

**Supplementary Information**

**Figures and legends**

**Figure S1. Example of a NeatSeq-Flow workflow:**

To demonstrate NeatSeq-Flow usage, we present a simple workflow (WF) operated on three samples, each with two FASTQ files of paired-end reads (Forward and Reverse). The example WF performs quality testing and trimming of the raw reads, alignment ("mapping") of the reads to a reference genome using two different programs and generation of sorted bam files. Additionally, the example WF creates a summary report on the statistics and quality of the raw reads, the trimmed reads and the mapping process.

Details of the example WF are as follows: The merge module copies the raw files from their original locations to the data directory, decompresses them, if necessary, and merges split files into a single file (per direction, in case of paired end reads). The fastqc_html and trimmo modules perform quality control with FastQC (<http://www.bioinformatics.babraham.ac.uk/projects/fastqc/>) and trimming with trimmomatic (1), respectively. Then, the sequences are aligned to the reference genome with Bowtie2 (2) (bowtie2_mapper module) and with BWA (3) (bwa_mapper module). The SAM files produced from these mapping steps are sorted and compressed with samtools (4). Finally, the results of the FastQC and mapping steps are graphically presented in a single MultiQC (<http://multiqc.info/>) report.

A graphical representation of the WF is shown in **A**. Note the two parallel mapping branches, by Bowtie2 and BWA, and the convergence of the two FastQC and two Samtools steps to the MultiQC step. **B** and **C** show the sample and parameter files for this WF, respectively. **D** shows the NeatSeq-Flow real time monitor. NeatSeq-Flow output directory structure, and the scripts produced for the example WF are presented in **E**.

**A tutorial of how to run this WF is available at NeatSeq-Flow Documentation:**

<http://neatseq-flow.readthedocs.io/en/latest/Example_WF.html>

**A.** **Graphical presentation:**

The diagram shown below is produced by an R script which is automatically generated in each run of NeatSeq-Flow. The diagram enables the users to visualize their own WF design, immediately after execution of NeatSeq-Flow script generator (see **E7,** "diagrammer.R" below). Each ellipse represents a step in the WF and is colored by module. The step name is written within the circle, and below it, in parentheses, is the module name. In the presented WF, there are two instances of the fastqc_html module, one testing the original FASTQ files and the other testing the files produced by the trimmo module. After the Trimmomatic step, there are two main branches, each running a different mapping program on the same samples (BWA and Bowtie2). Prior to mapping, a program-specific index building step is performed (BWA_Index_Builder and Bwt2_Index_Builder). Results from the quality testing and mapping steps are summarized in a graphical report produced by MultiQC (QC_and_Map_MultQC step). To achieve this, the QC_and_Map_MultQC step requires that all steps which produce input for it, namely all steps using the fastqc_html and samtools modules, will be executed before it. This is represented in the graph by the convergence of the four steps Fastqc_Merge, FastQC_Trimmomatic, Samtools_BWA and Samtools_bwt2 into the QC_and_Map_MultiQC step.

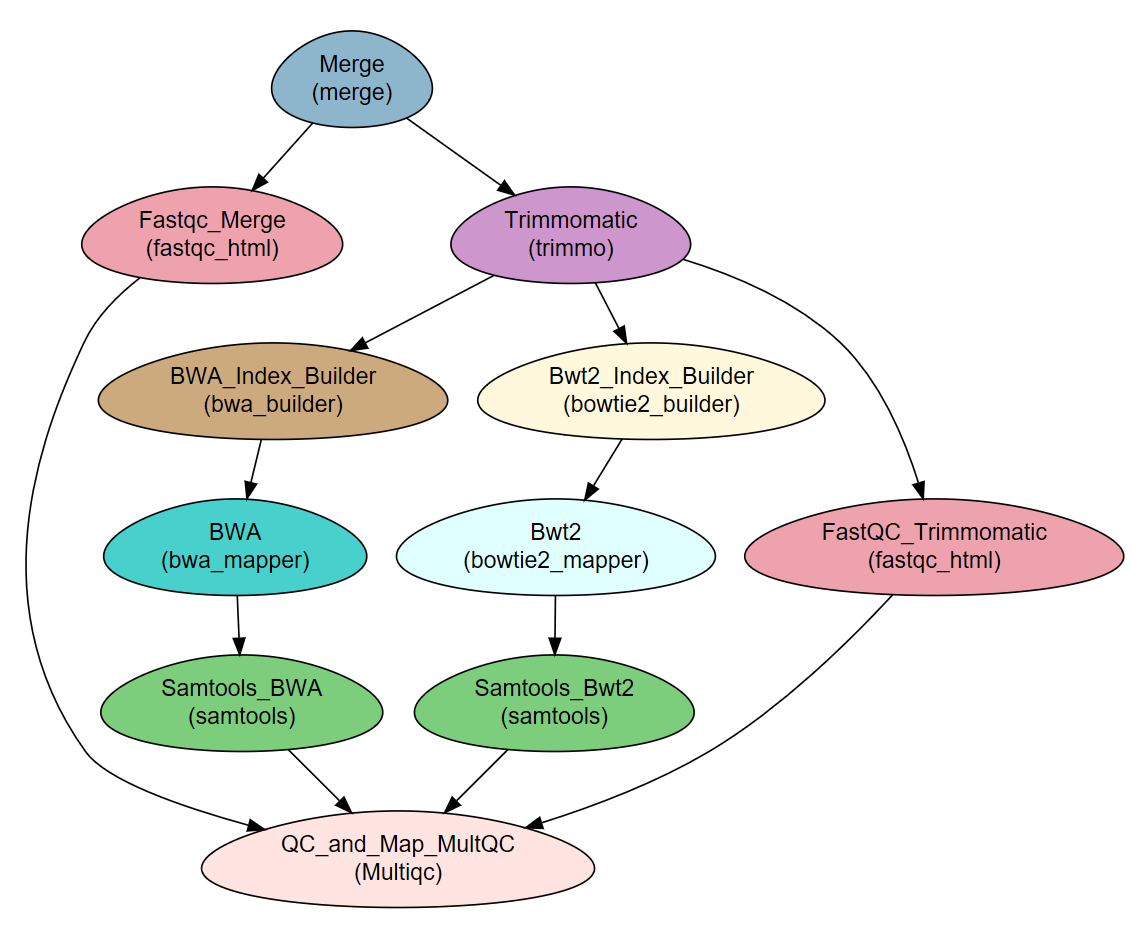

**B. Sample file:**

The sample file contains two sections: project-level raw files and sample-level raw files. The project-level section specifies raw files that are available to all samples (e.g. a reference genome). The sample level section specifies the samples’ raw files. Each raw file is described in one line, with file type (e.g. Single, Forward, Reverse, Nucleotide etc.), file path and sample ID (for sample level files) separated by tabs.

```
Title	Example_WF_From_the_manuscript

#Type 	Path
Nucleotide	Reference_genome.fasta

#SampleID	Type			Path
Sample1	Forward		Sample1F.fastq
Sample1	Reverse		Sample1R.fastq
Sample2	Forward		Sample2F.fastq
Sample2	Reverse		Sample2R.fastq
Sample3	Forward		Sample3F.fastq
Sample3	Reverse		Sample3R.fastq
```

Project level

Sample level

**C.** **Parameter file:**

The parameter file, written in YAML format, contains the WF design information and is composed of three main parts: "Global_params","Vars" and "Step_params".

In the "Global_params" section the user defines the global parameters of the WF such as cluster job scheduler parameters and the script-generator general parameters.

The "Vars" section contains the information that the user needs to update for the WF to operate in its own environment such as important files/directories and program executable paths. NeatSeq-Flow supports the use of CONDA environments to ease it’s installation on the user’s computer, including required HTS analysis programs, which are also included in the search path (for more information about using NeatSeq-Flow with CONDA see NeatSeq-Flow documentation, <http://neatseq-flow.readthedocs.io/en/latest/Example_WF.html#using-conda-virtual-environments>).

The "Step_params" section contains the actual design of the WF in the form of step definitions. For each step the user specifies the step unique name, module to use, step dependencies ("base") and step parameters. The step parameters can be divided into three types: 1. Module parameters that are module specific (e.g. the "mod" parameter set to "mem" in the BWA step). 2. Program's parameters such as the "script_path" parameter (the basic command to run the program) and the "redirects" parameters which are passed directly to the program command. 3. Step specific cluster parameters ("qsub_params") which indicate the step requirements from the cluster (e.g. the amount of CPU/memory needed for each job in this step).

The first step in any WF is always the merge module step. All other steps must contain the "base" parameter, which specifies previous step(s) that need to be completed before the current step starts.

More information about the parameter file’s format and full specification are available at NeatSeq-Flow Documentation (<http://neatseq-flow.readthedocs.io/en/latest/02.build_WF.html?highlight=parameter%20file#parameter-file-definition>)

Example parameter files for running with (1) and without (2) CONDA are shown below.

1. An example parameter file for running with CONDA:

```
## Global params:
Global_params:
    Qsub_opts: STR
    Qsub_path: /PATH_TO_YOUR_QSUB_Executable/
    Qsub_q: your.q
    Default_wait: INT
    module_path: PATH TO ADDITIONAL MODULES DIRECTORY
   # For running using CONDA environments:
    conda:
        path: {Vars.conda.base}
        env: {Vars.conda.env}

## Vars:
## Definition of variables used in this parameter file:
# Use the values stored in this section as follows:
   # {Vars.Programs.program1} refers to the PATH of program1
Vars:
    # For running using CONDA environments:
    conda:
        # The next line can be empty if running ‘export CONDA_BASE=$(conda info --root)’ from the
        # command line
        base: PATH TO CONDA BASE INSTALLATION DIRECTORY
        env: NAME OF THE CONDA ENVIRONMENT TO USE

    # Set the Parallel programming environment (PE) to instantiate in this workflow.
    Parallel_Environment:
        pe_name: STR
    Programs:
        FastQC: fastqc
        Trimmomatic:
            Bin: trimmomatic
            Adapters: PATH TO ADAPTERS FILE
        BWA: bwa
        bowtie2: bowtie2
        bowtie2_builder: bowtie2-build
        samtools: samtools
        multiqc: multiqc

## Step params:
## Workflow step and parameters definition:
## Definitions have the following form:
## Step NAME
##     parameter name: value
## USED MODULES IN THIS FILE:
    ## merge: Copies the raw files, decompresses zipped files and concatenates multiple files.
    ## fastqc_html: Runs the quality checking software ‘FastQC’ on all fastq files.
    ## trimmo: Trims the reads by quality using ' Trimmomatic ' program.
    ## bwa_builder: Create an index to be used by the BWA mapper.
    ## BWA_mapper: Maps fastq files to genomes using ‘BWA’ program.
    ## bowtie2_builder: Create an index to be used by the bowtie2 mapper.
    ## bowtie2_mapper: Maps fastq files to genomes using ‘bowtie2’ program.
    ## Samtools: Runs various ‘samtools’ on the SAM file produced by alignment modules.
    ## Multiqc: Creates a report for various programs.
## For information about the possible parameters for each module see the modules documentations.
## [ http://neatseq-flow.readthedocs.io/projects/neatseq-flow-modules/en/latest/ ]

Step_params:

    Merge:
       module: merge
        script_path:

    Fastqc_Merge:
        module: fastqc_html
        base: Merge
        script_path: {Vars.Programs.FastQC}
        qsub_params:
            -pe: '{Vars.Parallel_Environment.pe_name} 15’
        redirects:
            --threads: 15

    Trimmomatic:
        module: trimmo
        base: Merge
        script_path: {Vars.Programs.Trimmomatic.Bin}
        qsub_params:
            -pe: '{Vars.Parallel_Environment.pe_name} 20’
        todo: 'ILLUMINACLIP:{Vars.Programs.Trimmomatic.Adapters}:2:30:10 LEADING:20 TRAILING:20
               SLIDINGWINDOW:4:15 MINLEN:36'
        redirects:
            -threads: 20

    FastQC_Trimmomatic:
        module: fastqc_html
        base: Trimmomatic
        script_path: {Vars.Programs.FastQC}
        qsub_params:
            -pe: '{Vars.Parallel_Environment.pe_name} 10’
        redirects:
            --threads: 10

    BWA_Index_Builder:
        module: bwa_builder
        base: Trimmomatic
        script_path: '{Vars.Programs.BWA} index'
        scope: project

    BWA:
        module: bwa_mapper
        base: BWA_Index_Builder
        script_path: {Vars.Programs.BWA}
        scope: project
        mod: mem
        qsub_params:
            -pe: '{Vars.Parallel_Environment.pe_name} 20’
        redirects:
            -t: 20
            -B: 5

    Bwt2_Index_Builder:
        module: bowtie2_builder
        base: Trimmomatic
        script_path: {Vars.Programs.bowtie2_builder}
        scope: project

    Bwt2:
        module: bowtie2_mapper
        base: Bwt2_Index_Builder
        script_path: {Vars.Programs.bowtie2}
        scope: project
        qsub_params:
            -pe: '{Vars.Parallel_Environment.pe_name} 20’
        get_map_log:
        get_stderr:
        redirects:
            --end-to-end:
            -p: 20
            -q:

    Samtools_BWA:
        module: samtools
        base: BWA
        script_path: {Vars.Programs.samtools}
        qsub_params:
            -pe: '{Vars.Parallel_Environment.pe_name} 20’
        view: -buh
        sort: -@ 20
        flagstat:
        idxstats:
        index:
        stats: --remove-dups
        del_sam:
        del_unsorted:

    Samtools_Bwt2:
        module: samtools
        base: Bwt2
        script_path: {Vars.Programs.samtools}
        qsub_params:
            -pe: '{Vars.Parallel_Environment.pe_name} 20’
        view: -buh
        sort: -@ 20
        flagstat:
        idxstats:
        index:
        stats: --remove-dups
        del_sam:
        del_unsorted:

    QC_and_Map_MultQC:
        module: Multiqc
        base:
        - Fastqc_Merge
        - FastQC_Trimmomatic
        - Samtools_Bwt2
        - Samtools_BWA
        script_path: {Vars.Programs.multiqc}
```

2. The "Global params" and "Vars" sections in the example parameter file for running **without** CONDA (the "Step params" section is the same as in 1 above):

```
## Global params:
Global_params:
    Qsub_opts: STR
    Qsub_path: /PATH_TO_YOUR_QSUB_Executable/
    Qsub_q: your.q
    Default_wait: INT
    module_path: PATH TO ADDITIONAL MODULES DIRECTORY

## Vars:
## Definition of variables used in this parameter file:
# Use the values stored in this section as follows:
   # {Vars.Programs.program1} refers to the PATH of program1
Vars:
    # Set the Parallel programming environment (PE) to instantiate in this workflow.
    Parallel_Environment:
        pe_name: STR
    Programs:
        FastQC:          /FULL_PATH_TO/fastqc_Executable
        Trimmomatic:
            Bin:         /FULL_PATH_TO/trimmomatic_Executable
            Adaptors:    /FULL_PATH_TO/Adaptors_FILE
        BWA:             /FULL_PATH_TO/bwa_Executable
        bowtie2:         /FULL_PATH_TO/bowtie2_Executable
        bowtie2_builder: /FULL_PATH_TO/bowtie2-build_Executable
        samtools:        /FULL_PATH_TO/samtools_Executable
        multiqc:         /FULL_PATH_TO/multiqc_Executable
```

**D. Terminal Monitor**

NeatSeq-Flow real time monitor can be run to view the progress of the WF execution as well as previous or parallel WFs runs. Additionally, the monitor can indicate the occurrence of errors during the WF execution and therefore facilitates debugging. For each run of NeatSeq-Flow, the script generator assigns a unique Worfkflow ID. This ID will appear in the log file’s name of that run and accordingly may be used to monitor the run’s progression.

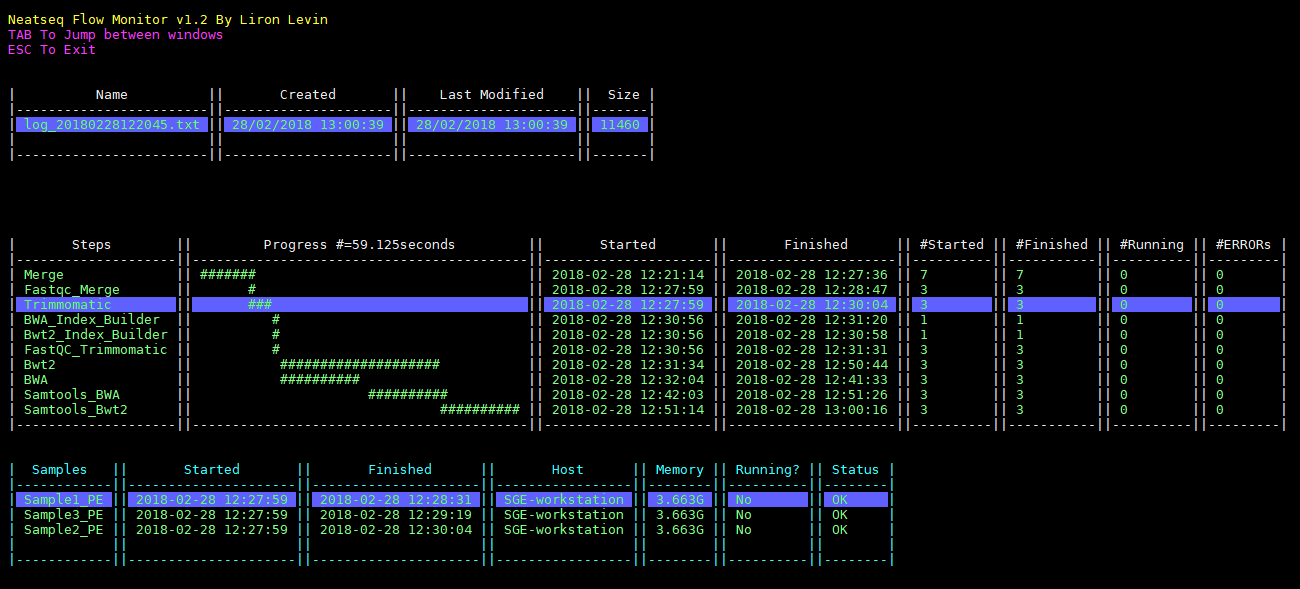

**E. Output directory structure:**

1. The main directory structure:

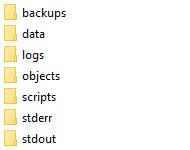

2. The **scripts** directory:
   The ‘00.workflow.commands.csh’ script (marked in yellow below) is the "master script" which executes the entire WF by calling the step-level scripts. The scripts beginning with consecutive numbers (e.g. ‘08.bowtie2_mapper_Bwt2.sh’) are step-level scripts, which execute entire steps by calling sample- or project-level scripts. The sample- or project-level scripts, which are the actual scripts running each step per sample or on the entire project, are contained in the equivalent directories, e.g. ‘08.bowtie2_mapper_Bwt2/08.bowtie2_mapper_Bwt2_Sample1.sh’.

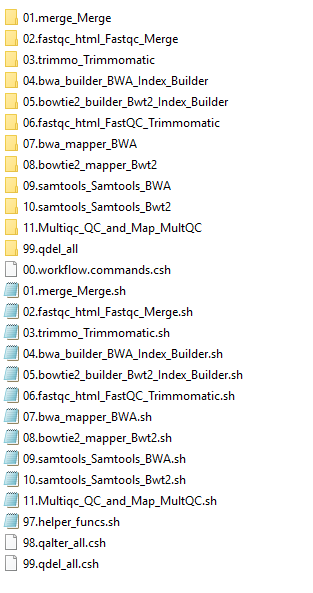

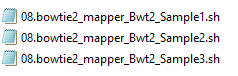

3. The **data** directory:
   In the data directory, the analysis outputs are organized by module, by step and by sample. Below is the data directory for the current example, showing the tree organization for the bowtie2_mapper and Multiqc modules.

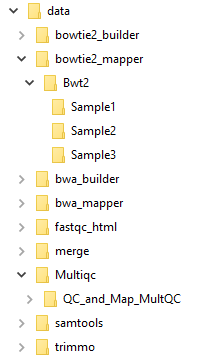

4. The **backup** directory:

The backup directory contains all used modules, as well as the sample and parameter files. As mentioned above, for each run of NeatSeq-Flow the script generator assigns a unique Worfkflow ID. This Worfkflow ID will appear in the name of the backup directory as well as in the backup parameter and sample file names of that specific run.

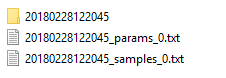

Worfkflow ID

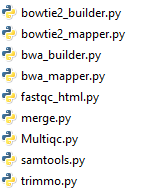

5. The **logs** directory:

The logs directory contains various logging files:

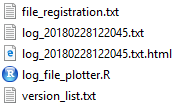

a. version_list.txt. A list of all the versions of the WF.
b. file_registration.txt. A list of the produced files, including md5 signatures, and the script and WF version that produced them.
c. log_file_plotter.R. An R script for producing a plot of the execution times (receives the name of the log file to plot as a single argument)
d. log_<workflow_ID>.txt. Log of the execution times of the script per workflow version ID.
e. log_<workflow_ID>.txt.html. Graphical representation of the progress of the WF execution, as produced by the log_file_plotter.R script.

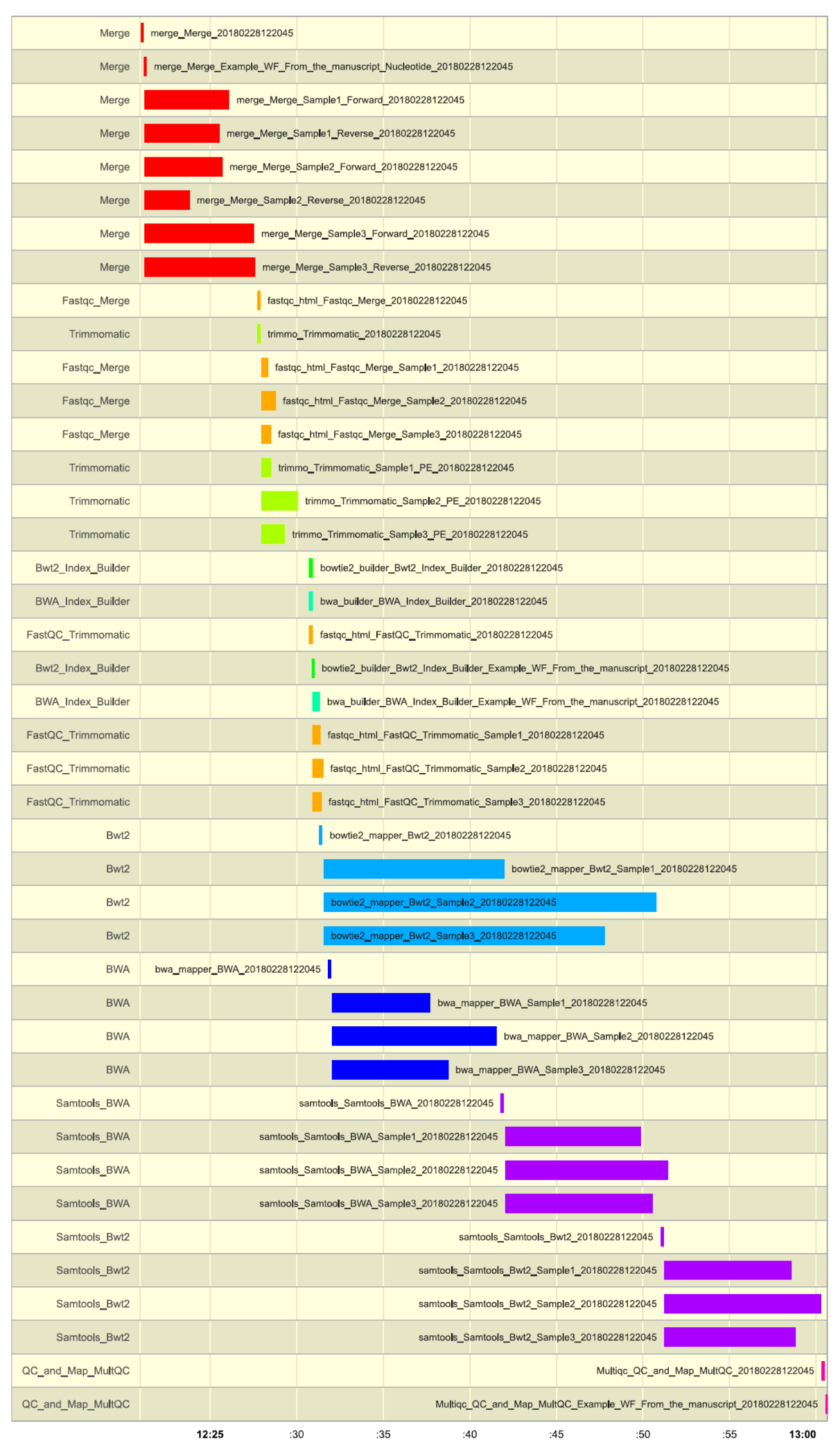

6. **stderr** and **stdout** directories:

The stderr and stdout directories store the script standard error and standard output, respectively.
These are stored in files whose names contain the module name, step name, sample name, workflow ID and cluster job ID.

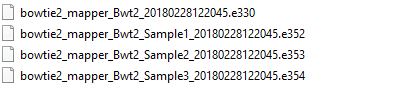

7. The **objects** directory:

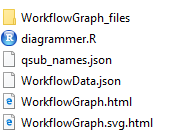

The objects directory contains various files describing the WF: An SVG diagram of the pipeline (WorkflowGraph.svg.html), an R script - diagrammer.R – for producing a DiagrammeR diagram of the WF (WorkflowGraph.html), and WorkflowData.json, containing all the WF data in JSON format, for uploading to JSON compliant databases etc.
The diagrammer.R script requires installing the 'DiagrammeR' and 'htmlwidgets' packages.

**Figure S2. Templates for new modules:**

New modules may be developed using templates, requiring only basic knowledge of Python. The templates can be obtained with the following links:

1. [A template for modules that create a script for each sample](https://neatseq-flow.readthedocs.io/en/latest/_downloads/NeatSeqFlow_ModuleTemplate_Sample.py).
2. [A template for modules that create one script for the entire project](https://neatseq-flow.readthedocs.io/en/latest/_downloads/NeatSeqFlow_ModuleTemplate_Project.py).

**Figure S3. Implementation of managing file transfer between steps:**

NeatSeq-Flow script generator maintains an internal data structure for passing file information between the different steps of the WF. The data structure contains two sections, samples and project. In the samples section, for each sample the available file type’s locations are stored and are indicated by circles; in the project section the file type’s locations that are available for all samples are stored and are indicated by stars; the colors of the shapes represent the different file types. Each step in the WF can access the data structure for its input file type locations and pass it to the next step with its output file type locations added. The scope of the step (sample level or project level) determines whether the step operates for each sample individually or for all samples together. In addition, it indicates to which section of the data structure the output will be added.

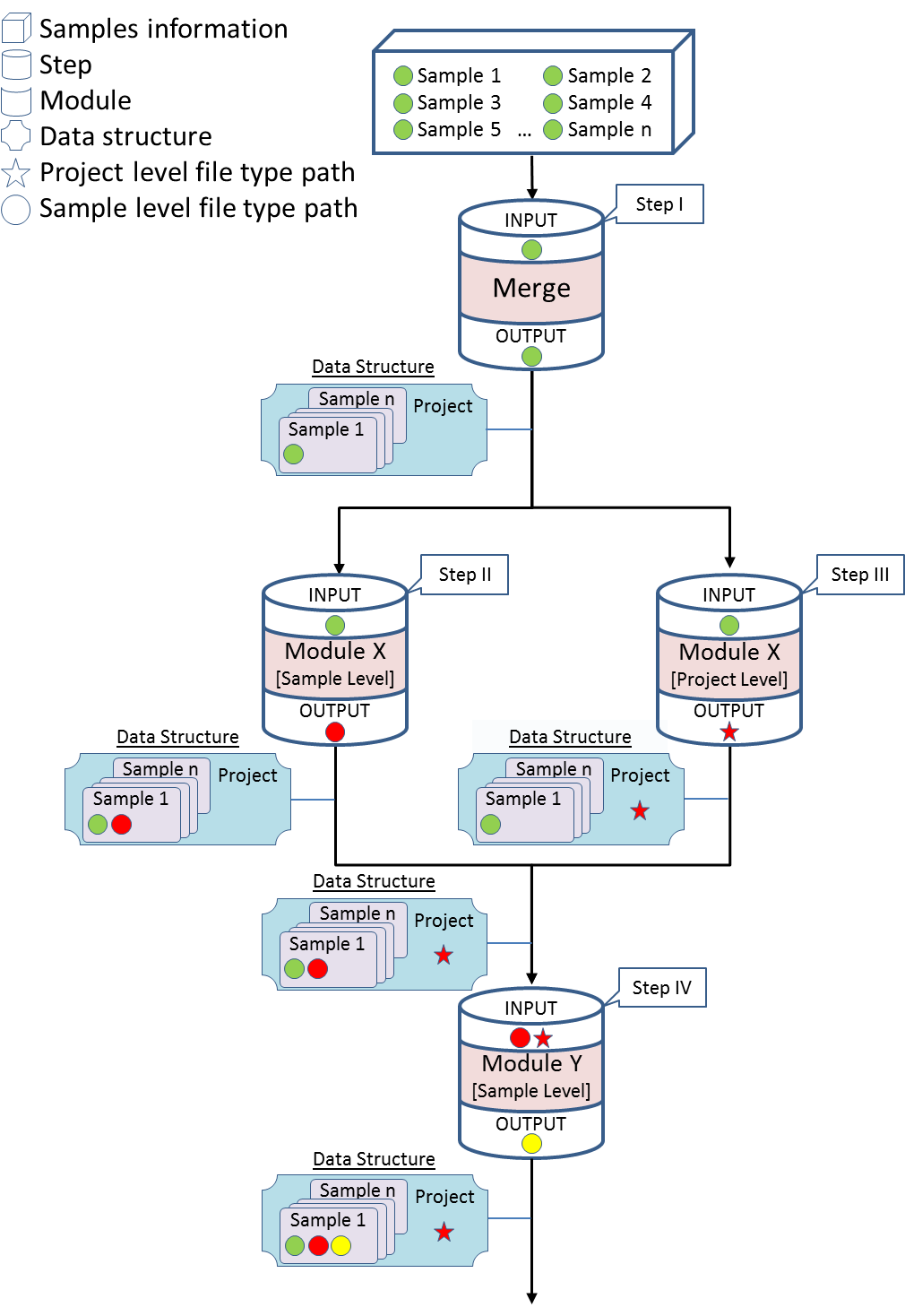

**Figure S4. Example of usage and implementation of the generic module:**

In the following example, a generic module is used to generate a BLAST database for each sample and a subsequent generic step queries each database with sequences from an external FASTA file. This example is a typical use of BLAST in many biological scenarios such as searching for virulence/resistance genes (whose sequences are in the external FASTA file) in bacterial genomes (the *de novo* assembly of reads from each sample). The Figure shows the fraction of the parameter file (in YAML format) on the left and the implementation using the data structure on the right.

**A.** Calling a generic module to generate a BLAST database (using makeblastdb) from each sample. This step can be used after (base:) any step that creates a nucleotide FASTA file (File_Type: fasta.nucl), e.g. after merge (if the raw files are in nucleotide FASTA format) or after a *de novo* assembly step. The location of the BLAST database for each sample is saved as a blast_db file type (File_Type: blast_db) for downstream use. **B.** Calling a generic module which performs a BLAST search (tblastn) of an external query protein fasta file (-query : path to query protein fasta file) against the previously generated BLAST database per sample. This step can be used after the Make_BLAST_DB step (base: Make_BLAST_DB). The user can pass additional parameters directly to the used program in the redirects section (e.g. –dbtype, –evalue, -num_descriptions etc.).

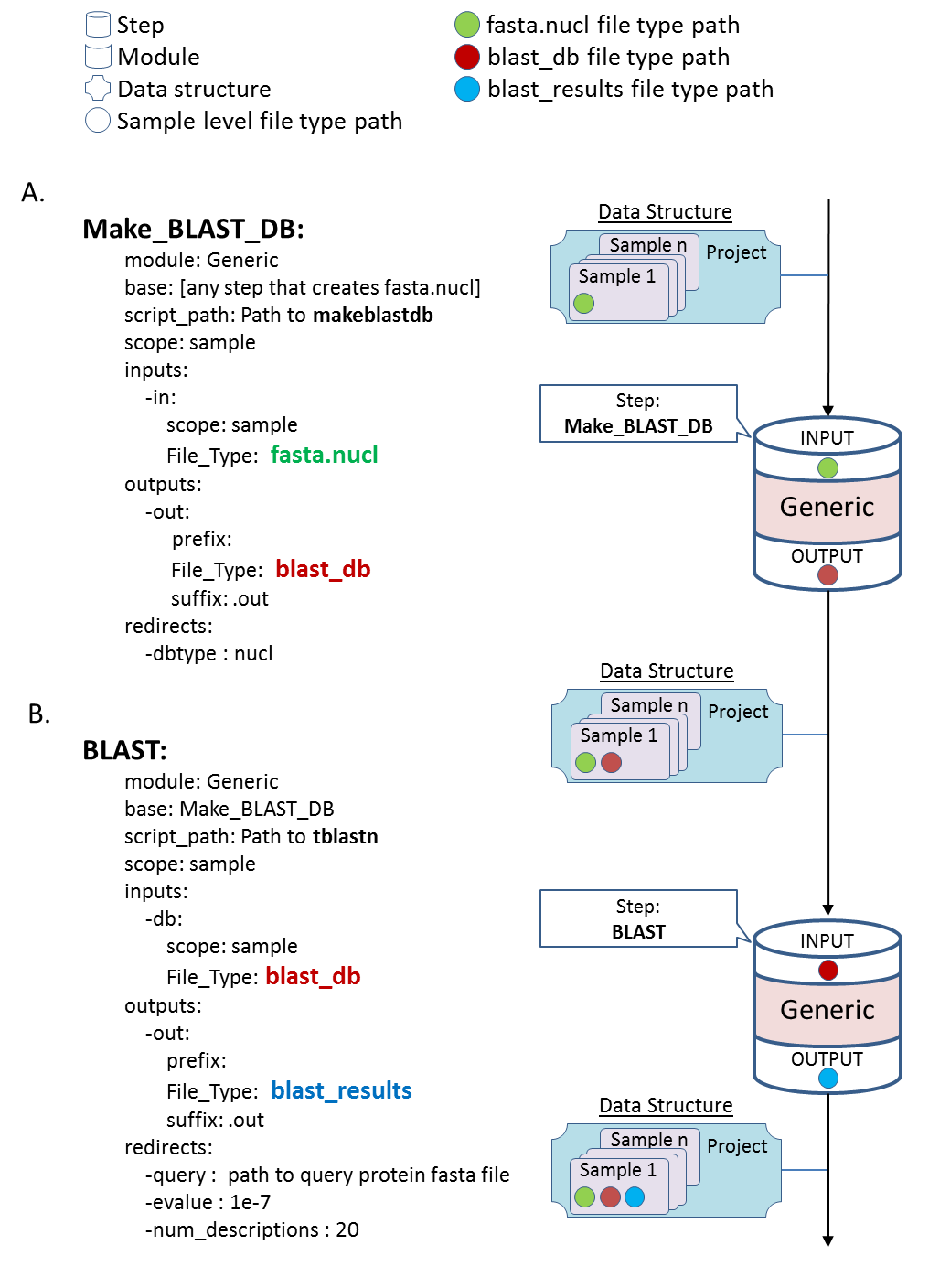
